# High-Throughput Conjugation Reveals Strain Specific Recombination Patterns Enabling Precise Trait Mapping in Escherichia coli

**DOI:** 10.1101/2025.02.26.640286

**Authors:** Thibault Corneloup, Juliette Bellengier, Mélanie Magnan, Arsh Chavan, Benoit Gachet, Zoya Dixit, Coralie Pintard, Alexandra Baron, Doreen Toko, Amaury Lambert, Alaksh Choudhury, Olivier Tenaillon

## Abstract

Genetic exchange is a cornerstone of evolutionary biology and genomics, driving adaptation and enabling the identification of genetic determinants underlying phenotypic traits. In *Escherichia coli*, horizontal gene transfer via conjugation and transduction not only promotes diversification and adaptation but has also been instrumental in mapping genetic traits. However, the dynamics and variability of bacterial recombination remain poorly understood, particularly concerning the patterns of recombined DNA fragments. To elucidate these patterns and simultaneously develop a tool for trait mapping, we designed a high-throughput conjugation method to generate recombinant libraries. Recombination profiles were inferred through whole-genome sequencing of individual clones and populations after selection of a marker from the donor strain in the recipient. This analysis revealed an extraordinary range of recombined fragment sizes, spanning less than ten kilobases to over a megabase—a pattern that varied across the three tested strains. Mathematical modelling indicated that this diversity in recombined fragment size enables precise identification of selected loci following genetic crosses. Consistently, population sequencing pinpointed a selected marker at kilobase-scale accuracy, offering a robust tool for identifying subtle genetic determinants that could include point mutations in core genes. These findings challenge the conventional view that conjugation always transfers large fragments, suggesting that even short recombined segments, traditionally attributed to transduction, may originate from conjugation.

## Introduction

Genetic exchanges occupy a pivotal position in genomics, serving two complementary roles. Firstly, by severing the physical linkage between alleles during a population’s evolutionary journey, it ensures that natural selection enforces meritocracy along the genome: alleles rise or fall on their own selective merit rather than being buoyed or hindered by their chromosomal neighbours (1). Secondly, this unshackling of allele effects from the surrounding genome is also central to geneticists’ ability to uncover the genomic underpinnings of phenotypic traits. From tracing genealogies to conducting quantitative trait loci (QTL) analyses and genome-wide association studies (GWAS), genetic exchange is crucial for pinpointing chromosomal regions linked to traits.

In bacterial genomes, where reproduction is uncoupled from genetic exchange, these transfers remain no less crucial. Processes such as conjugation, transduction, and natural transformation drive both genome evolution and scientific discovery (2). These mechanisms facilitate the spread of beneficial alleles or gene clusters, irrespective of their genomic background. Take the High Pathogenicity Island (HPI) in *Escherichia coli (E. coli)*: this genomic region, essential for efficient iron acquisition in diverse host environments, owes its dissemination within the species to homologous recombination following conjugation (3). Such transfers have not only fostered bacterial adaptation but have also enabled HPI identification as a major virulence determinant using agnostic statistical approaches such as GWAS (4). In addition, similarly to eukaryotic crosses, conjugation was instrumental in crafting the first bacterial genetic maps by tracking the inheritance of selectable markers (5).

While the mechanics of eukaryotic recombination are well-studied, bacterial recombination remains however more enigmatic. Unlike the single crossovers typical of eukaryotic meiosis, bacterial exchanges require double crossovers, resulting in varying integration lengths (6,7). Complicating matters further, as incoming DNA is often a threat, bacteria have evolved core stress responses or accumulated accessory defence systems (8), including restriction-modification genes, to degrade incoming DNA. Such defences fragment donor DNA, with the pattern of fragmentation potentially depending on strain genomic content. To better investigate these issues, we have developed a model of conjugation to unravel how conjugation shapes genome evolution and enables functional genomics.

Horizontal gene transfer in *E. coli* drives genomic diversification via two primary mechanisms. One involves phages erroneously packaging host DNA—a process known as transduction—while the other relies on plasmid-mediated conjugation. The relative importance of these mechanisms in nature remains contested. Comparative analyses of closely related genomes suggest that transduction predominates, with recombinant fragments spanning tens of kilobases, matching experimental observations (9,10). By contrast, conjugation transfers fragments potentially exceeding hundreds of kilobases. Mastery of these two processes has underpinned genetic engineering for decades. Transduction, with its low efficiency, has been most useful for transferring selectable markers across genetic backgrounds. Conjugation, by contrast, has been a workhorse for genetic mapping (5) and, more recently, for assembling complex genome constructs and high-throughput marker transfers (11).

Despite these advances, identifying the genetic basis of phenotypic traits in bacteria remains challenging. In a versatile species like *E. coli*, phenotypic diversity can stem from gene presence or absence, as exemplified by the HPI (3) or antibiotic resistance genes. Yet, over a decade of experimental evolution, coupled with whole genome sequencing, revealed that single-point mutations in conserved genes or regulatory regions can drive significant adaptive changes (12–14). Adaptation may even involve mutations in essential genes (15). Classical tools like transposon mutagenesis, while invaluable for gene inactivation studies, often fail to pinpoint these causative alleles. QTL-style crosses could address this limitation, provided the resulting genetic linkage is minimal.

In the present study, we developed a high-throughput conjugation method to enable large-scale genetic exchanges in *E. coli*. Our findings revealed that conjugation under identical conditions yields fragment lengths spanning more than two orders of magnitude, with recombination patterns and fragment lengths being strain-specific. This heterogeneity enabled precise identification of selected loci following crosses, a result corroborated by mathematical modelling. By shedding light on the extent of bacterial recombination, our approach not only provides a powerful tool for functional genomics but also highlights the potentially broader significance of conjugation in shaping *E. coli* genome evolution.

## Results

### 1. Rational for the design of a library of High-frequency Recombination (Hfr) variants

For more than 70 years, it has been known that the transfer of chromosomal fragments through conjugation can occur at high frequency with a conjugative plasmid inserted in the chromosome (16). Parts of the chromosome directly adjacent to the integrated conjugative plasmid, 5’ upstream of the origin of transfer (*oriT*), are transferred from the High-frequency recombination (Hfr) bacteria (17) (Fig 1A). As a consequence, conjugation from a single given Hfr to a recipient strain results in a biased transfer of genetic material proximal to the integration site. A major objective for exploiting conjugative recombination is to discover unknown loci imparting novel phenotypic traits in natural isolates of *E. coli* (18). Since the position of such loci is unknown, a biased transfer proximal to a fixed position in the chromosome may limit their discovery. Therefore, to increase the success of discovery, we need to initiate transfer from multiple positions along the chromosome simultaneously (Fig 1B). To achieve an unbiased and uniform transfer of DNA from a donor strain, rather than using a single Hfr, we decided to create a library of Hfr variants, each having the conjugative plasmid integrated at a different site along the chromosome.

**Figure 1.**
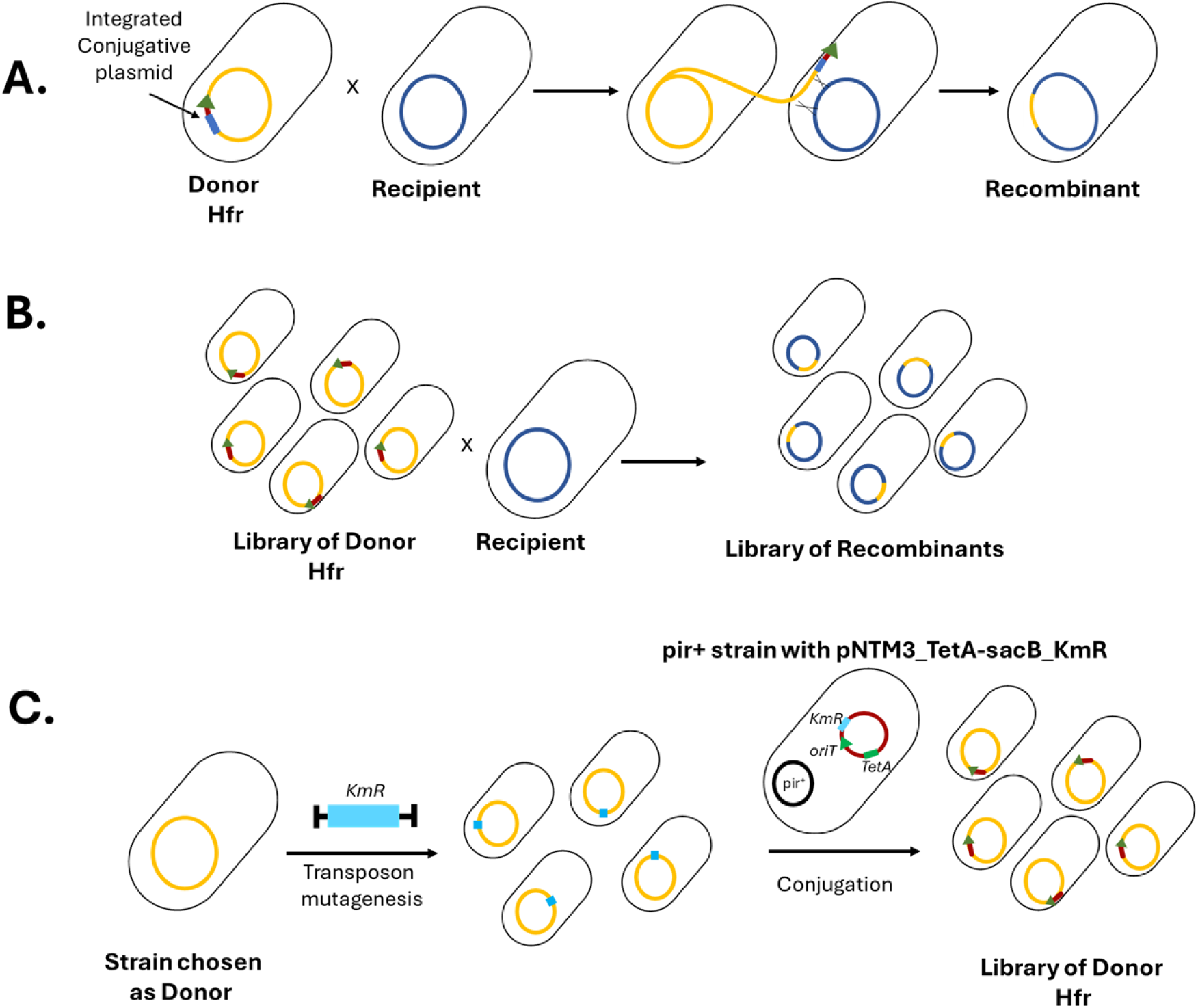
Creation of single locus Hfr donor versus multi-locus Hfr donor. A) Chromosomal DNA transfer from a single locus Hfr strain to a recipient showing position-dependent bias in the transferred DNA fragment. The green arrow and dark blue box represent the origin of transfer of the plasmid and for simplicity the integrated plasmid. B) DNA transfer between a library of Hfr donors and a single recipient. Here, different fragments are transferred between the donor and the recipient. C) A landing pad, here a Kanamycin cassette, can be integrated at a random position in the genome with a transposon mutagenesis approach based on Mariner or Tn5 transposons. Homology between the cassette on the plasmid and the one on the chromosome promotes integration of the plasmid in a different location in each mutant.

Conjugative plasmid integration occurs mostly through homologous recombination between plasmid and chromosome sequences that are highly similar (17). The similarity between the plasmid and the chromosome is often the result of insertion sequences (IS), which can easily jump across the genome (19). The distribution of IS is variable between strains, and most IS elements are present in a few copies in the chromosome (20). Therefore, IS offer only a limited, variable and biased number of insertion sites. We decided to use a conjugative plasmid that has been shortened to be IS-free: pNTM3 (11). This suicide plasmid was designed to enhance its integration into the chromosome by replacing its native origin of replication with an R6K origin of replication (Fig S1A). This origin requires the presence of the *pir* gene to maintain the plasmid (21). The *pir* gene is provided in trans in a permissive strain that allows replication and transfer of the plasmid. In strains lacking *pir*, the plasmid, encoding an antibiotic resistance gene, can be grown in media with the corresponding antibiotic only if the plasmid has integrated into the genome to maintain the resistance genes. The pNTM3 plasmid has been previously modified by the integration of certain chromosomal fragments to integrate at various sites on the chromosome (18). However, our goal was to be able to integrate it at a random position in the chromosome of various *E. coli* strains. For this purpose, we decided to introduce a genomic landing pad with homology to the pNTM3 plasmid. We took advantage of the well-characterized transposon mutagenesis technique (Fig 1C) (22). Using the machinery of a Tn5 transposable element (or Mariner see *Method*), we can easily introduce a Kanamycin resistance cassette (*KmR*) randomly into the chromosome of an *E. coli* strain. Although not all natural isolates are amenable to genetic manipulation, many strains can be modified/mutated using transposon mutagenesis. To use the Kanamycin landing pad for conjugal recombination, we introduced a Kanamycin resistance gene (*KmR*) into the plasmid near the origin of replication to generate plasmid pNTM3_TetA-sacB_KmR (Fig S1A). Therefore, upon transfer of this *KmR* pNTM3 plasmid within the transposon mutagenesis library, we expect to obtain a library of variants with this conjugative plasmid randomly integrated at different sites across the genome.

### 2. Heterogeneous integration rates of the conjugative machinery

We first tested whether we could obtain efficient and consistent rates of pNTM3_TetA-sacB_KmR plasmid integration into the chromosome via homologous recombination between kanamycin resistance genes. To this aim, we used strains from the Keio collection (23). Conveniently, this collection is composed of strains, each with the kanamycin resistance gene inserted to replace and inactivate all non-essential *E. coli* genes. We realized a transfer and integration with 14 strains with different inactivated genes across the *E. coli* genome and a negative control - BW25113 (Parent of the Keio collection, derivative of K12) - lacking the kanamycin resistance gene (*KmR*). First, we observed a low frequency of integration in the negative control. This suggests residual integration independent of the presence of our landing pad. Second, there was considerable heterogeneity in plasmid integration rates across genomic loci. While we failed to have integration at some positions such as *nfrA*, we obtained integration rates of up to 10^5^ CFU counts HFR/Pntm3 donor, in others, such as *ybeX* (Fig 2B). The integration rates were however overall low, below 10^−8^ per donor, and sometimes similar to those of the negative control suggesting that the Hfr libraries produced will have biased integration sites.

**Figure 2.**
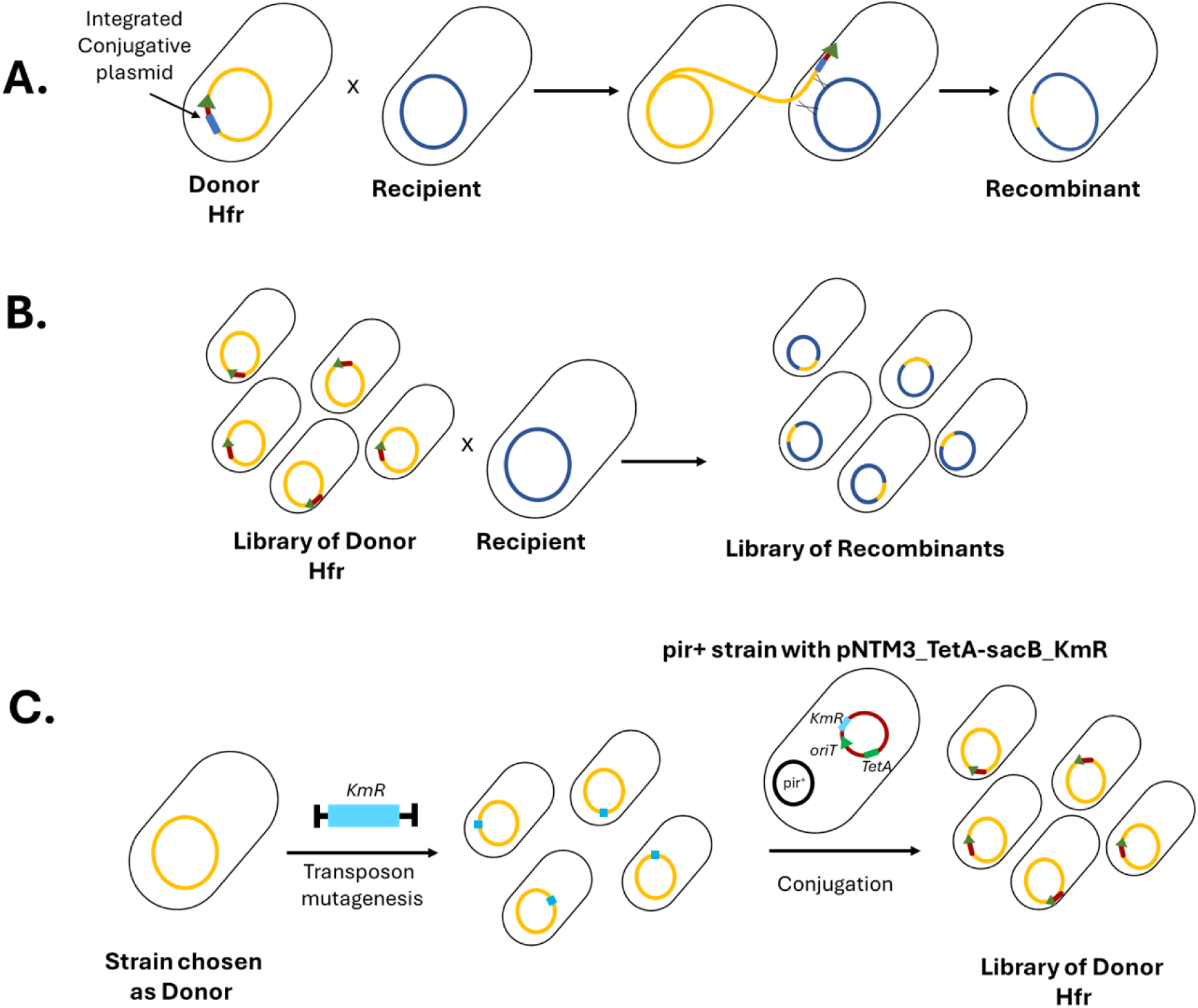
Position-dependent variability in Hfr construction. A) Schematic representation of the pNTM3_TetA-sacB_KmR plasmid with tetracyclin resistance (*tetA*) and *KmR* as the homologous marker for integration on the recipient genome. The pNTM3_TetA-sacB_KmR host is a diaminopimelic acid (DAP) auxotroph and harbours the *pir* gene to replicate the R6K origin of replication. Hfr cells were grown on LB + tetracycline without DAP to select for the recombined Hfr recipients. B) Hfr frequency is measured as the log10 ratio of CFU per mL of trans-conjugant recombinants (selected on LB + tetracycline) to CFU per mL of donors (selected on LB + DAP + tetracycline). We represented CFU counts from 3 independent replicates

### 3. Inducing targeted integration with double-strand break

To overcome the integration bias of pNTM3_TetA-sacB_KmR in the chromosome, we decided to improve recombination by using Cas9-mediated targeted DNA double-strand break (DSB) and Lambda-red recombination (24). The Cas9 endonuclease is directed by a guide RNA (gRNA) for targeted DNA double-stranded break (DSB). The gRNA contains a spacer region with a 20 bp homology to the target site, which is proximal to a 5’-NGG-3’ Protospacer Adjacent Motif or PAM. Cas9:gRNA interaction with the PAM and gRNA::DNA binding at the homologous target site induces the DNA DSB. In *E. coli*, efficient Cas9-mediated targeted DNA DSB is cytotoxic and results in cell death. Cells with a repair template to replace the PAM or the spacer using phage-mediated homologous recombination gain immunity to the targeted DNA DSB. Therefore, Cas9-mediated DNA DSB can select for precise genome modifications coupled with PAM/spacer mutations introduced using phage-mediated recombination (24). In addition, recent studies have shown that having a targeted DNA DSB can also improve the efficiency of recombination (25–28).

We induced a DNA DSB in genomic Kanamycin landing pad using the Cas9 endonuclease from the *Streptococcus pyogenes* CRISPR system (29). We tested repair using three different gRNAs. The cytotoxicity of the killing efficiency can vary between gRNA and the editing efficiency is correlated to the killing efficiency. Therefore, we first tested the gRNA killing efficiency in Keio strains with an integrated copy of the kanamycin resistance gene. All had similar efficiencies, so we chose one (sgRNA1) and introduced it into plasmid pNTM3_TetA-sacB_KmR (Fig S1A). To rescue the cells from the targeted DNA DSB using recombination, we modified the plasmid kanamycin resistance gene to replace the corresponding PAM motif with a synonymous PAM mutation (SPM). Therefore, genomic integration of the plasmid would replace the chromosomal *KmR* gene with the *KmR* gene with SPM and provide immunity from the Cas9-mediated DSB to select for the plasmid integration. The resulting plasmid was named pNTM3_sgRNA.

To promote integration of pNTM3_sgRNA, we added the plasmid encoding Cas9 and the Lambda-red recombination system to recipient cells by electroporation. We induced the system to facilitate recombination between the sequences flanking the cut site in the chromosome and the repair template present on the pNTM3_sgRNA plasmid (see methods) (30).

Indeed, with the Cas9/recombineering-mediated insertion protocol, we achieved a much higher integration efficiency (Fig S1B) and less heterogeneity between sites, with a rate greater than 10^−5^ CFU counts of Hfr per donor for all but one locus (13 out of 14) (Fig 3B). Therefore, this two-plasmid method significantly improved pNTM3 recombination efficiency to construct an Hfr at different loci of interest.

**Figure 3.**
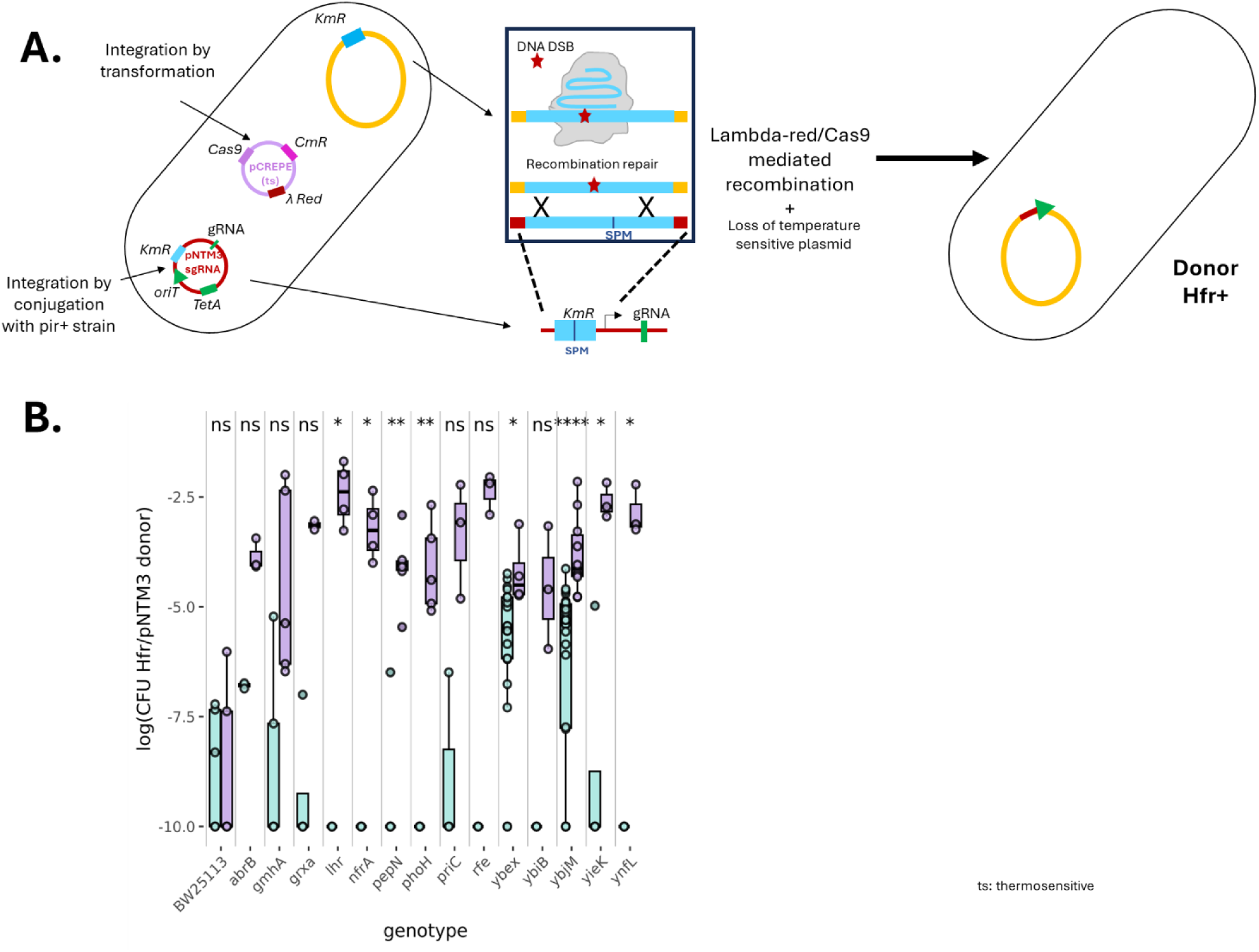
Cas9-Lambda-red recombination-mediated Hfr construction. A) Schematic representation of the gRNA and repair template for Cas9-Lambda-red recombination-mediated Hfr construction in the recipient. B) Log10 of the Hfr production rate with the Cas9-Lambda-red approach (purple) across the 14 loci and a control compared to the production without induced double-strand break (cyan). Wilcoxon test was used for statistical analysis. ns : non-significant, * : p value ≤ 0.05, ** : p value ≤ 0.01.

### 4. Validating the ability of constructed Hfr to transfer the *galK* marker

To validate that the selected strain had the plasmid chromosomally integrated and was a functional Hfr, we tested their ability to transfer DNA to a recipient strain. The Hfr strains constructed, using the selected variants from the Keio collection, had the plasmid integrated at variable distances from the *galK* gene. We therefore tested each Hfr’ ability to transfer the functional *galK* gene into a recipient strain modified to have a non-functional *galK* gene. The protein GalK (galactokinase) is required for growth on galactose as the sole carbon source. Using CRISPR-Cas9, we added multiple stop codons to inactivate the *galK* gene preventing any possibility of reversion by mutations. Consequently, in this strain, only recombination of a fully functional gene upon conjugative transfer from the Hfr donor could restore the *galK* function (Fig 4A) (31). We then crossed each constructed Hfr with the *galK-* recipient. Upon conjugation, we found that the strain at 100kb could transfer the *galK+* allele with extremely high efficiency. Corroborating previous observations, we observed that integration at a higher distance resulted in lower transfer rates (from 939 kb on, the transfer rates were below detection limit), but the strain with an Hfr at 300 kb could still transfer the *galK*+ allele with 20% efficiency (Fig 4B).

**Figure 4.**
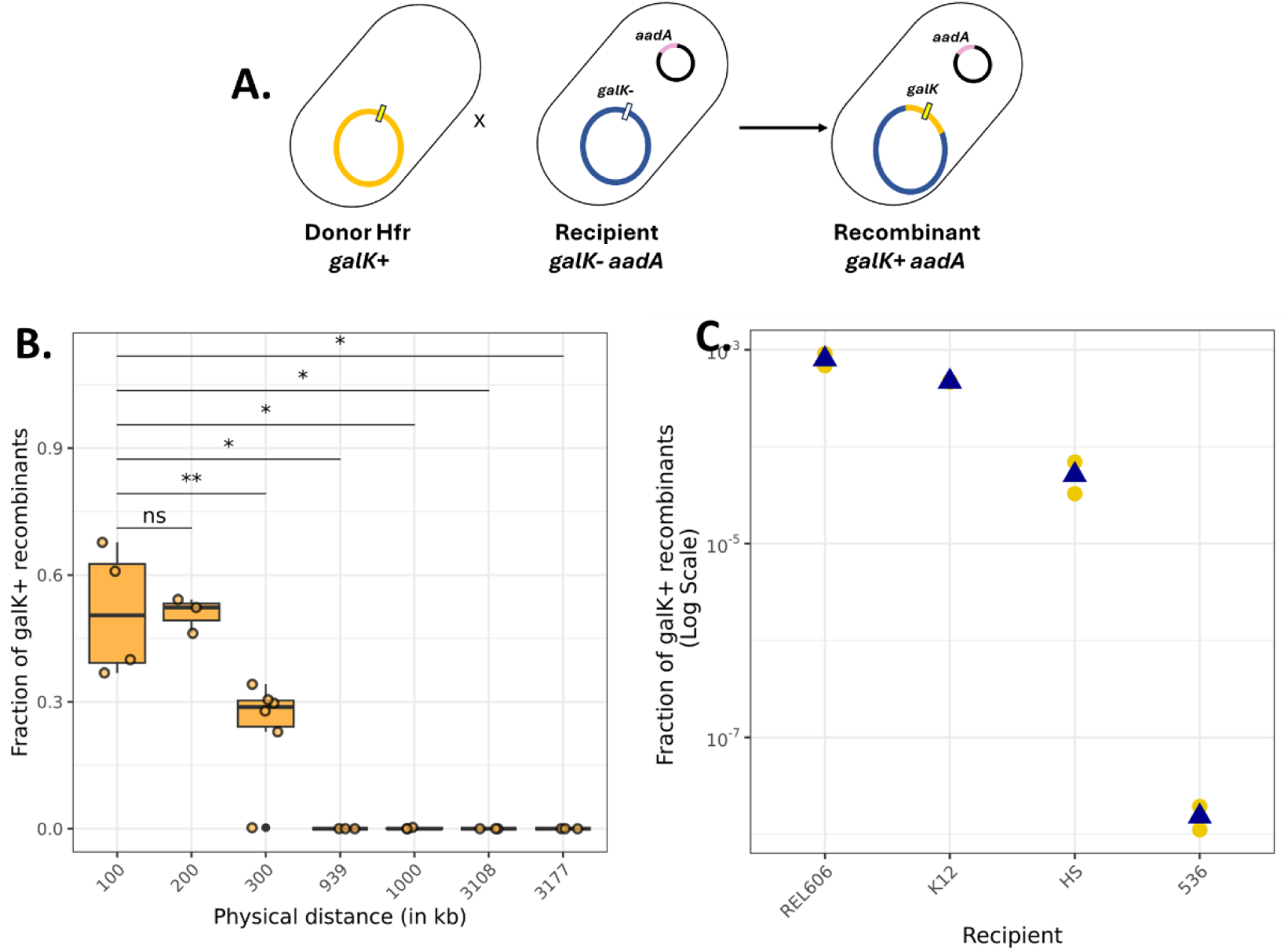
Testing for conjugation. A) A schematic representing crosses between *galK*+ Hfr donors and *galK*-spectinomycin recipients to select transconjugant recombinants of M9 minimal media with galactose and spectinomycin. B) Conjugation efficiency measured as the ratio of CFU per mL of trans-conjugant recombinant recipients (selected on M9 + galactose + spectinomycin) to CFU per mL of all recipients (selected on M9 + glucose + spectinomycin) for Hfr constructed at different distances from the *galK* locus. Statistical comparisons were made using the Wilcoxon test, ns: non-significant, *: p-value ≤ 0.05, **: p-value ≤ 0.01. C) Conjugation efficiency measured as the log10 ratio of CFU per mL of trans-conjugant recombinant (selected on M9 + galactose + spectinomycin) to CFU per mL of all recipients (selected on M9 + glucose + spectinomycin) for Hfr library crossed with *galK*-spectinomycin recipients in different genetic backgrounds *E. coli* K-12 BW25113 (control), REL606, HS, and 536. Mean are represented by blue triangles and data points by yellow dots.

### 5. Creating and testing a library Hfr

After functional validation of our targeted Hfr constructs, we tested if we could create a library of Hfr. We used for that a transposon library in which the *KmR* landing pad is randomly inserted in the chromosome. Transposon libraries were made in the genetic background of the laboratory strainsBW25113 and REL606. We then used the afore mentioned Cas9/recombineering-mediated system to integrate the pNTM3_sgRNA plasmid across the different sites of integration of the transposon. This protocol led to the selection of 10^3^ to 10^4^ colonies that were pooled in Hfr libraries.

To test if in these Hfr libraries, the integration of pNTM3_sgRNA conjugative plasmid occurred across multiple loci along the genome, we investigated the integration sites. We adapted a method for transposon sequencing with our libraries of Hfr donors to find the precise locations of the landing pad in which pNTM3_sgRNA was integrated (32). With this approach, we only amplify a short DNA sequence that spans the extremity of the transposon and the proximal chromosomal sequence (Fig S2A) where the integration has occurred. We then used sequencing to extract the exact insertion positions and observed pNTM3_sgRNA plasmid was integrated at multiple loci across the chromosome.

The genome architecture may lead to biases in the integration of the plasmid. One source of bias is differences in copy number variation between different regions of the genome (with higher copies closer to the origin of replication, compared to the terminus) due to differences between rate of chromosomal replication and cell division. In addition, organization of the *E. coli* chromosome into macrodomains may result in a biased accessibility of their DNA (33). We therefore investigated the spread of insertion sites along the chromosome using the genomic coverage derived from transposon sequencing data (the number of reads that match a specific insertion site) (Fig S2B). We compared the density of insertions along the chromosome as the log fold-change in coverage compared to the coverage of the *Ter* macrodomain. There are however two ways to analyse the data: we can focus on the insertion sites (red in Supp. Fig S2B), or the coverage of these insertion sites (blue on Fig S2B). Insertion sites refer to the identified positions of insertions and likely refer to independent insertions during the process of Hfr construction. Coverage of these sites corresponds to potential differences in the representation of the various Hfr strains in the library, but also integrates potential biases associated with sequencing (34). As mentioned, higher copy number of the *Ori* macrodomain within the cells leads usually to higher coverage of that region relative to the *Ter* macrodomain in sequencing data.

Analysing density profiles along the genomes with bins of 250kb, we observed that, there was a bias for both metrics: fewer integration sites were found in the *Ter* macrodomain compared to the rest of the genome (Fig S2C) and the coverage of insertions in the *Ter* macrodomain was lower. The bias was visible using both metrics, but was, as expected, more marked considering coverage of insertion sites (up to 10-fold, Fig S2C) than density of insertion sites (up to 3 fold, Fig S2C). This is not unexpected given the afore mentioned *Ori* to *Ter* coverage bias observed in whole genome sequencing. The biases in the density of insertion sites suggest a biological bias during transposon mutagenesis, that likely results from the lower number of copies and/or lower accessibility of the *Ter* macrodomain. Despite this bias, integrations of the conjugation machinery were spread along the chromosome and our library could therefore be used to transfer any locus in the genome to a recipient strain.

To test if this library could transfer DNA to various genetic backgrounds, we realized conjugation between a donor K12 BW25113 Hfr library (constructed twice independently) and 4 recipients with different genetic backgrounds: the laboratory strains BW25113 and REL606, the latter being the ancestor of the famous experimental evolution initiated by Richard Lenski (35), the natural isolate HS (36), and the pathogenic strain 536 (37). While the first three correspond to strains from phylogroup A of *E. coli*, the latter from B2 phylogroup is more genetically distant. All four strains were made *galK-* and resistant to spectinomycin via the presence of a plasmid. After the mating with the Hfr library, we plated on minimal media with galactose and spectinomycin to select the recombinants and get rid of donors and recipients. The number of recombinants varied largely depending on the recipient’s genetic background (Fig 4C). We estimated the recombination efficiency as the percentage of number of recombinant colonies (selected by plating minimal media with galactose and spectinomycin) to the number of total recipient colonies (estimated by plating on non-selective minimal media with glucose and spectinomycin). In BW25113 and REL606 backgrounds, the recombination efficiency was up to 5-10 %, This efficiency reduced to ∼0.6% in HS giving only a few colonies of recombinants and it goes as low as 0.0001% when 536 was the recipient. We could explain the lower recombination efficiency by the fact that strain 536 is expressing toxins thus killing the donor strains during the conjugation assay (Fig S4).

### 6. Analysis of the genome of the recombinants

Since we initiated the transfers from multiple loci, we expected to observe different patterns of recombination along the genome (Fig 5A). For the various crosses we made, we sampled 20 clones for sequencing (only 8 in the case of 536 due to low efficiency). Because parental genomes are known, we could sort the reads from the sequencing as sequences specific to the donor genome or the recipient or common to both. Here, reads specific to one strain are reads that match preferentially to one genome due to polymorphism differentiating the two stains. In this analysis, we only kept reads that matched both genomes and discarded the one matching exclusively a single one. Our aim was to focus on the backbone genome conserved between the two strains and not on the parts specific to one or the other strain. We then trained a hidden Markov chain model to delineate transitions between two states: matching stretches for either the donor or the recipient. As illustrated in Fig 5B-E, all genomes but one had a clear recombined region around the selected site.

**Figure 5.**
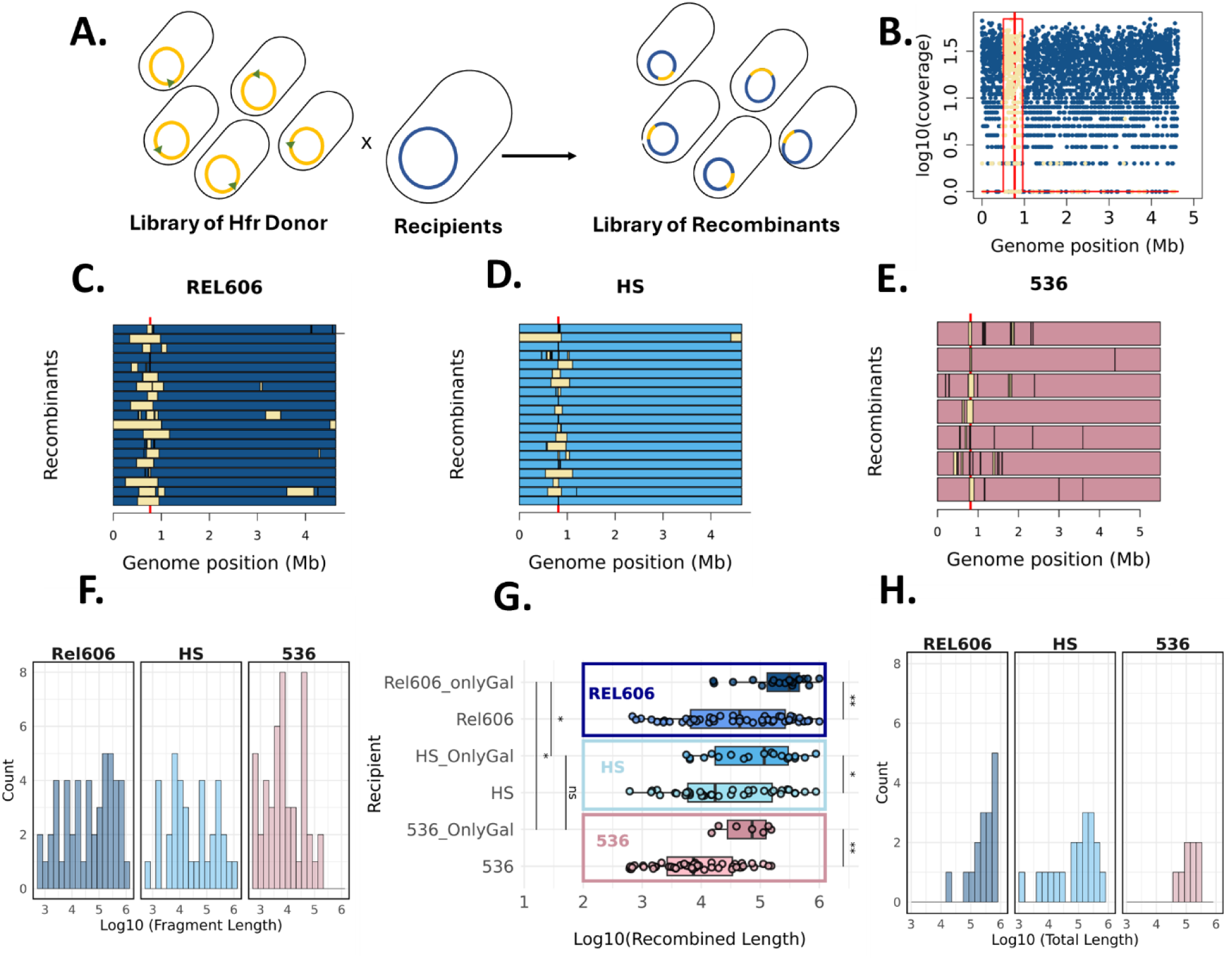
Genome sequencing of recombinant clones. A) A schematic representing varied hybrids from conjugation between a donor Hfr library and a recipient. B) Logarithm 10 of the per kb coverage of the genome with reads specifically mapping to the recipient genome (dark blue) or the donor one (yellow). Red line shows the *galK* loci position. Red broken line shows the delineation of the recombined fragment using Hidden Markov Models (HMM). C, D, E) Recombined fragments (yellow fraction) across the genomes for recipient REL606 (C), HS (D) and 536 (E). Red line indicates the *galK* loci position. F) Distribution of recombination fragment length (log10 scale), colours correspond to the different recipients (REL606, dark blue, HS, light blue, 536 light pink). G) Boxplot showing the size of recombination fragments for each recipient (log10 scale), for fragments including the selected marker (upper box) and the ones not including it (lower box, lighter colour). Wilcoxon-test was used for statistical analysis. ns : non-significant, * : p value ≤ 0.05, ** : p value ≤ 0.01. H) Distribution of the total length recombined (log10) per genome of recombinants (REL606, dark blue, HS, light blue, 536 light pink).

The patterns of recombination were however quite different between recombinants and between recipient strains (Fig 5C, D, E). First, many strains showed a simple fragment of recombination that overlapped the *galK* but the length of this fragment was very variable: from a few kilobases (1.5 kb or 7.5 kb) or more, up to a full megabase with a fragment as long as 1.103 Mb. Transfer size can therefore span about 3 orders of magnitude. Second, many strains harboured not only a recombined fragment around the locus of selection *galK*, but also several other recombined regions in the neighbourhood. This is compatible with an incoming DNA fragment being cut and recombined into pieces (7). This pattern was most drastic with strain 536 as recipient. Interestingly, we noticed that a few recombinants revealed other distant distinct sites of recombination. Finally, a few strains had not resolved the recombination in the colony that was sequenced as the recombined regions appeared to have both overlapping donor and recipient-specific sequences.

Overall, the distribution of the recombined fragments varied across strains (Fig 5C-G). The median size of recombined fragments was 184,487.3 bp, 115,776.5 bp and 22,638 bp for the recipients REL606, HS and 536 respectively (Fig 5F, G) while when we looked at the fragments that included the *galK* locus, their lengths were significantly greater across all strains (324,598.8, 194,828.6 and 78,631.86 bp median size in REL606, HS and 536 respectively) (Fig 5G).

We also tested the lengths’ differences between the *galK-*containing fragments across recipient strains and found that there was a significant difference between REL606 recombinants compared to HS and 536 recombinants (Fig 5G). When we looked at the total length of recombined material, the difference between strains remained but the distribution peaked around 200,000 to 400,000 bp (corresponding to 5.3 to 5,6 in log10 base) (Fig 5H).

### 7. Expected power to detect a selected site

Sequencing of individual clones confirmed the occurrence of recombination and provided critical insights into the length, position, and number of recombined fragments. Beyond this descriptive analysis, the experiment can be viewed as a form of cross coupled with selection, which enables the detection of loci under selection in a QTL-like framework. However, unlike the genetic exchanges typical of eukaryotic systems used in QTL approaches, the recombined fragment lengths here are highly variable. This raises the question of how such variability impacts the ability to pinpoint loci under selection.

To explore this generically, we employed mathematical modelling. We analyzed the statistical properties of recombined fragments conserved among a pool of selected recombinants near the allele under selection. The genomic region where donor DNA is consistently found across all selected recombinants is likely to contain the selected allele—here, the functional *galK* allele. Recombined fragments were modelled as independent, identically distributed intervals with positions uniformly distributed across the genome. This model demonstrated that for a large number (n) of selected recombinants, the length of donor DNA shared by all recombinants scales with the harmonic mean of the length of the recombined fragment encompassing the selected loci divided by the number of recombinants. Specifically, the expected size of the region likely to harbour the selected locus, denoted as, Λ_*n*_, is given by

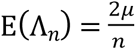, where 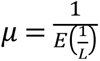 represents the harmonic mean of the recombined fragment lengths that include the selected loci (see Proposition 4.1 in the Supplementary Materials).

This result is significant because the harmonic mean is particularly sensitive to smaller values, meaning that the presence of short fragments—just a few kilobases in length—can substantially reduce the size of the shared recombined region across recombinants. Consequently, the model suggests that the observed heterogeneity in recombined fragment lengths may, counterintuitively, enhance the precision of locus detection under selection.

### 8. Analysis of the pool of recombinants

To test our ability to detect with precision the selected loci, we decided to use another approach based on sequencing a pool of recombinants. Using this methodology, we can analyse the whole population and look at the density of donor-specific reads along the genome to visualize patterns of recombination at the pool scale.

Because we are enforcing a strong selection at *galK* loci, we expected a clear and strong signal of donor DNA integrated into the recipient. For each of the recipients’ backgrounds, we identified a pick of donor-specific sequences around the *galK* loci (Fig 6A, B, C). The delineation of the target region was even more precise using the ratio of donor to recipient-specific reads along bins of 1 kb along the genome. The maximum of that ratio landed exactly in the *galK* gene for all crosses (Fig 6A, B, C). The pattern of decay of the donor DNA presence differed between the recipient strains (Fig 6D) with a decay less marked in REL606 than in HS and 536. This pattern is consistent with the distribution of recombination length detected as longer recombination tracks should lead to a slower decay.

**Figure 6.**
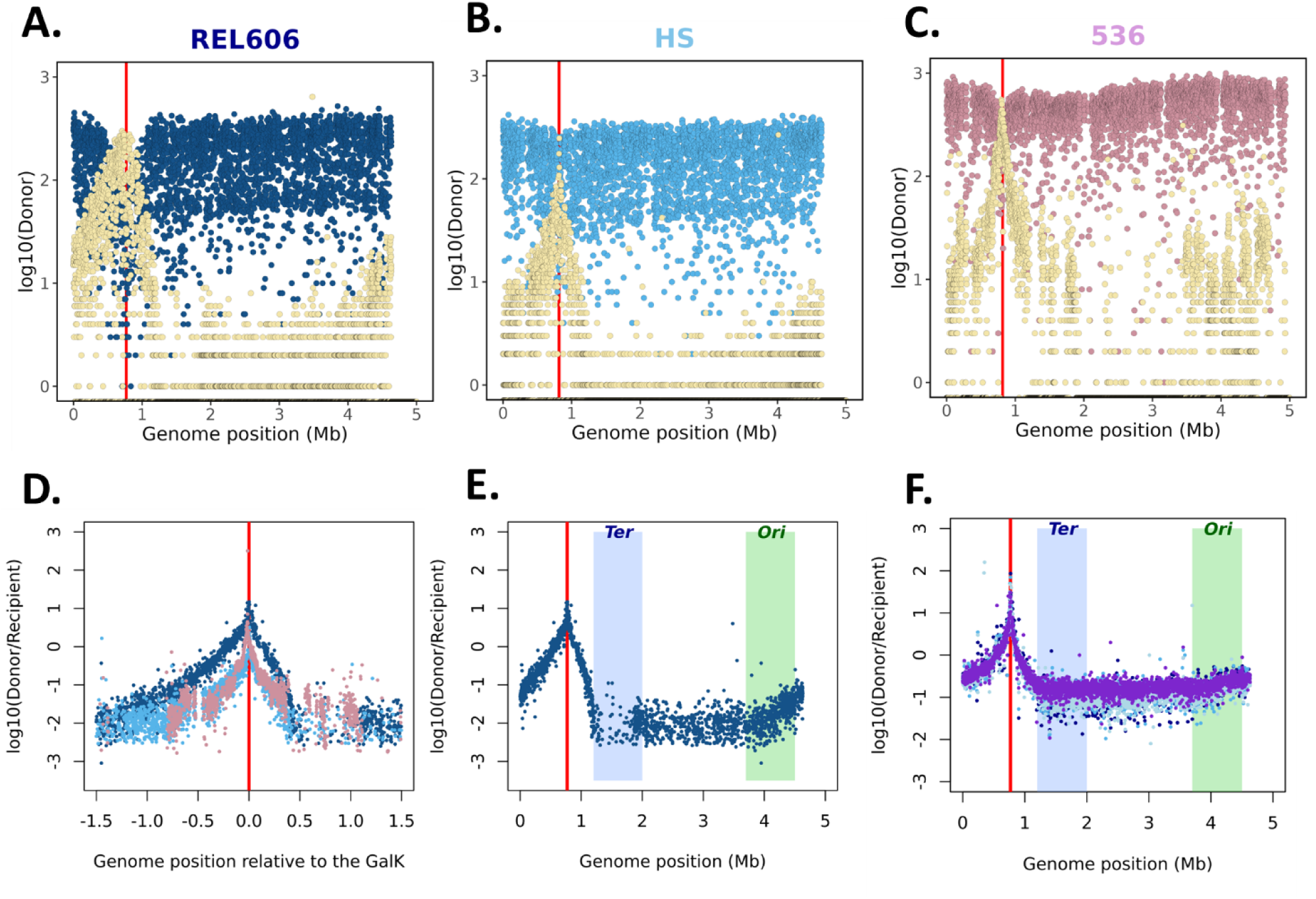
Pooled Genome sequencing. A, B, C) Log10 of the per kb coverage of the genome with reads specifically mapping to the recipient genome (in A dark blue for REL606, in B blue for HS, in C pink for 536) or the donor one (yellow). Red lines show the *galK* loci position. D) Log10-ratio of donor-to-recipient coverage along the genome relative to *galK* position for the three recipient genotypes (dark blue for REL606 blue for HS, red for 536.E) Log10-ratio of donor-to-recipient coverage along the genome for REL606 pool with the Terminus *Ter* (light blue) and *Ori* (green) macrodomain positions shown. F) Log10-ratio of donor-to-recipient coverage along the genome for two different genotypes Δ*matP galK-* and *galK-*. Red line shows the *galK* loci position with the Terminus *Ter* (light blue) and *Ori* (green) macrodomain positions are shown.

For all recipients, we observed a steeper reduction of donor DNA towards the *Ter* macrodomain (Fig 6E). This pattern could result from the integration bias of the conjugative plasmid in our library. However, we also know that the *Ter* macrodomain is structured and made less accessible through the interaction between the MatP protein and the *matS* sites that are frequent in that macrodomain (33). To test if that structuration of the *Ter* Macrodomain could reduce the efficiency of recombination and contribute to the observed pattern, we used a Δ*matP* mutant as a recipient. We observed no difference between Δ*matP* and wild type (Fig 6F) suggesting that recombination was not affected by macrodomain structural organization and that the biases resulted most likely from our skewed distribution of conjugative plasmid integration sites.

### 9. Selection at different loci and precision of the genomic analysis

We demonstrated through the *galK* crosses that we could extract critical insights into the recombination patterns in pools and the variability of recombined regions using individual clones. To verify that these observations were not artefacts stemming from the chosen selection locus or its chromosomal position, we generated a new library of REL606 recipients using an alternative marker for counter-selection. By employing transposon mutagenesis, we randomly inserted a cassette containing the toxin *tse2* under the control of a rhamnose-inducible promoter (38). Upon rhamnose induction, these mutants expressed the toxin and undergo cell death, enabling the selection of recombinants incorporating donor DNA near the insertion site, as the donor lacks the toxin. From this library, we selected three individual clones as recipient with Tse2 cassette inserted at distinct random positions and recombined them with a library of *E. coli* K-12 BW25113 donors. Genomic analysis of the resulting recombinant pool provided profiles of recombination events (Fig 7). By analyzing the ratio of donor- to recipient-specific reads, we observed that the selection pattern closely resembled the one seen with the *galK* crosses. Selection effects were highly localized and specific across all pools. Since recipients were now selected using a chromosomal marker (a zeocin resistance cassette introduced into the chromosome), donor DNA was counter-selected at the corresponding loci.

**Figure 7.**
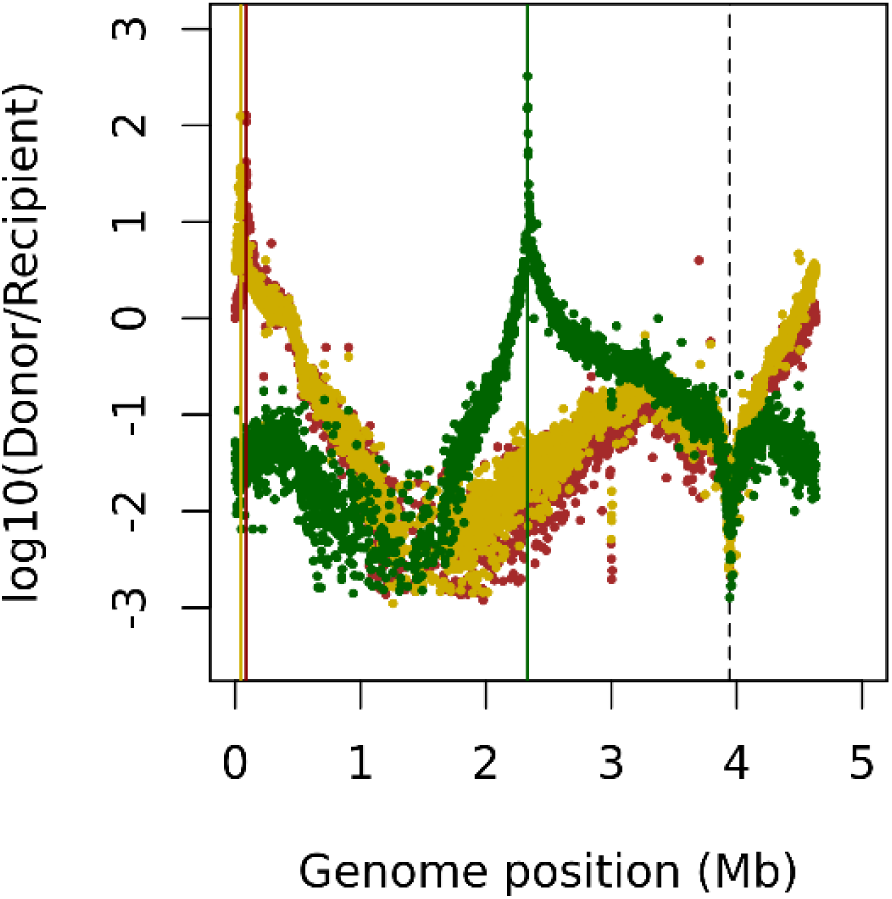
Pooled Genome sequencing of tse2 recombinants. Log10-ratio of donor-to-recipient coverage along the genome for the three REL606 recipients with different *tse2* positions (respectively in dark red, yellow, dark green) The vertical lines (red, yellow, dark green) correspond to the maximum of the log10 ratio Donor/Recipient for each of the sequenced pools (*tse2* position in the recipient). The black dotted line corresponds to the position of *ZeoR* present the recipient background and used for selection.

The next question was whether we could localize the position of these toxin insertion sites with the same precision as the one achieved with the *galK* loci, where three independent replicates yielded an accuracy of 1 to 1.5 kilobases. To address this, we analyzed the chromosomal position of the maxima in the log10 ratio of Donor/Recipient reads (Fig. 7) for each pool of recombinant. This peak serves as our best estimate of the location of the Tse2 cassette in the recipient strain. We then designed primers positioned 1 kb and 1.5 kb upstream and downstream of the inferred peak, along with primers specific to the Tse2 cassette itself. PCR amplification using the 1 kb primer set successfully detected the cassette in the inferred region for two out of the three samples. For the third sample, detection required pairing primers 1.5 kb from the inferred peak with those targeting the cassette (Fig S3A.). Thus, across all three samples, we identified the position of the locus under selection with an accuracy of ±1.5 kb. Therefore, we showed that our method could be used for efficient and precise QTL mapping.

## Discussion

In this study, we present a novel high-throughput approach for genomic conjugation. This method relies on the generation of a library of mutants called Hfr donor libraries, each mutant having a different chromosomal integration site for a conjugative plasmid. To promote random integration into the chromosome, we employed homologous recombination between the plasmid and a transposon cassette, the latter having been randomly integrated into the genome via transposon mutagenesis. To enhance integration rates and reduce biases, we resorted to CRISPR-Cas9 system to induce double-strand breaks in the chromosome, alongside the use of lambda Red recombination machinery to enhance recombination efficiency.

Before discussing the results we obtained, it is important to acknowledge several limitations of our protocol. One major limitation of our study lies in the use of a specific conjugation system while many different conjugative elements exist. Second, in the present study, we focussed so far on the selection at a specific marker though we showed the patterns were conserved at other sites. Another issue is that our library of donors is not homogeneously spread around the chromosome presumably due to reduced integration of the transposon and integration of plasmids in the *Ter* macrodomain near the terminus of replication. This prevents us from using our protocol to investigate the heterogeneity of recombination along the chromosome observed using comparative genomics (39). Despite these limitations, our protocol allowed us to successfully generate recombinant libraries from crosses between the K-12 laboratory strain and three different *E. coli* strains.

Genomic analysis of these recombinant libraries revealed an unexpected diversity in the lengths of donor fragments incorporated, ranging from a mere kilobase to over a million base pairs— significantly broader than previously reported ranges (7,9,10). Earlier genome-wide studies had suggested that recombined fragments were typically in the tens of kilobases, and therefore associated with bacteriophage-mediated transduction (10), while conjugation was linked to larger transfers involving several hundreds of kilobases. Our findings indicate that, even in tightly controlled laboratory settings, conjugation can yield extensive diversity in recombined fragments. This suggests that even short recombined fragments detected in genomic studies could originate from a conjugation event.

This observed variability in recombined fragment size has significant implications for the precise identification of loci under selection. By analysing the length variability and distribution of fragments, we can pinpoint genomic regions subjected to selection with 1 to 2 kilobases precision. Mathematical modelling suggests that when selecting for a specific allele from a donor strain, the size of shared fragments among recombinants correlates with the harmonic mean of the length of the recombined fragment encompassing the selected loci. Since the harmonic mean is sensitive to smaller values, the broad size range we observe enables precise identification of selected loci. This leads to two key consequences: first, it allows for the identification of causative alleles under strong selection; second, it suggests that conjugation may influence natural genome evolution in unforeseen ways.

Detecting the genetic source of a strain specificity remains an important challenge in microbiology. Within the same species, a large variability of phenotypes is observed including some traits of importance such as virulence or antibiotic resistance (40,41). For many of these traits, gene presence and absence are the major sources of diversity, and alternative methods including transposons mutagenesis have been used for decades with success. However, some important phenotypic diversity is explained by allelic variation within conserved regions of the genome rather than gene presence and absence as seen in experimental evolution (12,42). To discover these types of variations our approach may be relevant, though further investigation will be needed to test its efficiency when the locus studied is under moderate selection.

The considerable diversity in recombined segments challenges also the existing assumptions about genetic linkage and bacterial conjugation. While conjugation was supposed to involve large fragments, our results suggest that the size of the block transferred in full linkage is much lower than expected. This finding introduces an element of unexpected plasticity in bacterial genome evolution. The sharp decline in the fraction of donor DNA around selected loci indicates that following transfer and subsequent selection, genetic diversity within the population is largely preserved. This limited linkage disequilibrium may account for the mosaic nature of bacterial genomes and their rapid adaptability to diverse environments.

Our analysis also identified variations in recombination profiles among recipient strains, hinting at an inverse relationship between recombined fragment sizes and the phylogenetic distance between donor and recipient strains. Several factors could account for this variability. First, defence mechanisms, such as restriction-modification systems, might fragment incoming DNA, leading to lethal consequences for bacteriophages but allowing multiple integrations of homologous DNA (7). Additionally, sequence divergence may reduce the number of perfect homology sites necessary for recombination initiation, thereby hindering integration (38). Finally, selection may disfavour strains harbouring recombined fragments due to their disruptive effects on genomic organization. All these hypotheses have to be further investigated notably by applying our protocol to a large number of strains and selected mutants.

## Materials and methods

### 1. Growth Media

For bacterial growth, we used LB media or M9 minimal media. LB was purchased from Life Technologies (Thermofischer) (reference 12795027) with 5 g/L yeast extract, 10 g/L tryptone, and 10 g/L NaCl; 15 g/L agar added as needed. In each 100 mL M9 media, we added M9 salts (prepared using premixed powder from BD Biosciences Difco 5X (reference 248510)), 0.4% Glucose or Galactose or Rhamnose, 2 µM MgS0_4_, 0.1 µM CaCl_2_, and Thiamine 0.2 µg/mL. For plates, 1.6% Select agar was added. Recombination efficiencies were measured using Bacto McConkey agar with 0.4% galactose (15). When appropriate, antibiotics were added to the medium as follows: chloramphenicol (34 µg/ml), kanamycin (50 µg/ml), spectinomycin (50 µg/ml), gentamicin (15 µg/ml), tetracycline (10 µg/ml) or zeocin (50 µg/ml).

### 2. Strains, plasmids and cloning

To construct the high-frequency recombination (Hfr) donors, we used the donor strain BW38029 (*rrnB3 φ (lacZp4105(UV5)-lacY) 638 ΔlacZ hsdR514Δ(araBAD)567 rph+, ΔdapA::pir)*. These strains contain the suicide conjugative plasmid: pNTM3 (11). pNTM3_sgRNA was derived from pNTM3, an RP4 conjugative plasmid modified to have a *pir*-dependent R6K, a spectinomycin resistance cassette for selection. This plasmid is carried in a *pir+ recA-* host *ΔdapA::pir*, in which they replicate as plasmids but do not integrate into the chromosome. Upon transfer to a *pir*-*recA+ F-* strain, the plasmid integrates into the chromosome by homologous recombination, at a site determined by the homology region.

pNTM3 plasmid (Fig S1A.) was first modified by the introduction of the cassette containing a resistance gene to tetracycline and a gene conferring sucrose sensitivity-respectively *TetA* and *sacB* by recombineering (Datsenko-Wanner method) (43) using the following primers Pnmt3_aadA_tet_Wup and Pnmt3_oriR_sac_Wdn (primers sequences available in Supplementary Material). Using the same method, *KmR* from pKD4 was incorporated in this plasmid using the primers used TC_gRNABB_F and TC_gRNABB_R. Validation of integration used a set of common test primers k1 and k2. This plasmid, pNTM3_*TetA-saB_KmR* was first used to tests integration on the chromosome using clones from the Keio collection (Figure 2). Because of unexpected heterogeneity in integration patterns, we modified the plasmid further. An intermediate vector called pVec was then created by Gibson assembly using the backbone of a plasmid containing colE1 and *aadA* and a fragment of in pNTM3_*TetA-saB_KmR* containing *finO*, *KmR*, *TetA*, *sacB* and *R6K.* The following primers were used: Con_INS_F, Con_INS_Rev, Con_BB_for and Con_BB_Rev. To add a SPM on KmR, we used mutagenesis oligos (kan_SDM_R and kan_SDM_f) and ligation. Using Gibson Assembly, we then cloned a gRNA1 instead of *sacB* in pVec using the following primers TC_gRNA_INS_F and TC_gRNA_INS_R. The choice of gRNA (sequences available in the Supplementary) was realised by testing their efficiency using the primers kan_gRNA1_f, kan_gRNA1_r, kan_gRNA2_f, kan_gRNA2_r, kan_gRNA3_f and kan_gRNA3_r. Finally, a cassette from pVec containing *finO, KmR*, TetA* and sgRNA was introduced using Datsenko-Wanner recombineering into pNTM3. The final plasmid is denoted as pNTM3_sgRNA.

Individual donors to test the CRISPR-cas9-recombineering-mediated Hfr construction were constructed using the strain *E. coli* K12 BW25113 (*lacI^q^ rrnB_T14_ ΔlacZ_WJ16_ hsdR514 ΔaraBAD_AH33_ ΔrhaBAD_LD78_*) and its derivatives from the Keio collection. We disrupted several genes with a kanamycin resistance gene: *abrB, gmhA, lhr, nfrA, pepN, phoH, priC, rfe, ybeX, ybiB, ybjM, yieK, ynfL*.

For CRISPR cas9-recombineering-mediated genome modification, we used the pAM053 plasmids (43). The gRNA plasmid was expressed under the J23119 promoter using a plasmid purchased from Addgene (https://www.addgene.org/71656/). The *galK-* mutation was introduced by CRISPR-cas9-mediated recombineering using a construct named G22 which has been reported in a previous study (15). The kanamycin cassette for transposon mutagenesis was amplified using the pSAM_*Kan* plasmid. Cells were mainly grown on LB unless mentioned otherwise. All our conjugation experiments were performed using strains *E. coli* K-12 BW25113, *E. coli* REL606, *E. coli* HS, *E. coli* 536.

### 3. Lambda-red recombination and CRISPR-cas9-mediated recombineering

At least 100 ng of DNA and plasmid used for editing were dialyzed using a 0,025 µm membrane. The protocol for CRISPR-cas9-recombineering and Lambda-red recombineering were the same. We started in LB with Chloramphenicol overnight cultures of cells containing either a plasmid encoding Lambda-red recombination genes or a plasmid containing *cas9* and *lambda-red* recombination genes at 30°C. The next day, fresh 100 mL cultures were launched with a 100-fold dilution of the overnight cultures in fresh media and grown at 30°C up to a mid-log OD measured at 600 nm (OD600) of 0.4– 0.5. When the cells reached the desired OD, the flasks were placed in a shaking water bath set at 42°C for 15 minutes to induce the *lambda-red* recombination operon. After 15 minutes, the cells were immediately placed on ice and chilled for 15-20 minutes. The cells were then centrifuged at 7,500×g at 4°C for 3 minutes. The media was discarded and the pellet was resuspended in 25 mL ice-cold 10% glycerol and resuspended gently. The pellet washing step was repeated with 25 mL 10% glycerol 5 times and with 10 mL 10% glycerol one final time. The pellet was finally resuspended to a final volume of 300 µL in 10% glycerol. The cells were mixed with the dialyzed DNA and were introduced in the 1mm electroporation cuvettes. The cell-DNA mixture was electroporated at 1.8 kV and recovered using 1 mL of LB media in 1.5mL tubes. The cells were recovered for 2 to 3 hours at 30°C and then plated on LB agar plates with the required antibiotics.

The protocol for making competent cells was the same as above but with two differences: 1) There was no induction step. 2) The cells were grown at 37^°^C unless they harboured a temperature-sensitive plasmid.

### 4. Construction of the transposon library

The transposon library was prepared either using EZ-Tn5 transposase or by conjugation between recombination-proficient MDF *pir*+ cells with the plasmid pSAM_*Kan* (conjugation protocol above).

When using the hyperactive form of Tn5, we constructed our transposon library by mixing the EZ5-Tn5 transposase with the Kanamycin cassette to be inserted into the genome. The Kanamycin cassette was first amplified by PCR using pSAM_*Kan* plasmid as a template. The ME transposase recognition sequence was added using the primers : Ez_Tn5_Kan_for and Ez_Tn5_Kan_rev. The PCR product was extracted using gel purification and dialyzed using 0.025 µm membranes to remove excess salts. Subsequently, the DNA was phosphorylated using the NEB T4 polynucleotide kinase (PNK) enzyme by mixing 44 µL of the DNA with 5 µL of the T4 ligase buffer with ATP and 1 µL of the T4 PNK enzyme. The reaction was incubated at 37^°^C for 60 minutes and the product was PCR purified (Qiagen). The purified product was then mixed with the Ez-Tn5 transposase as: 1 µL of PCR product + 1 µL of 100% glycerol + 1 µL of ultrapure DNase free RNase free water + 1 µL of Ez-Tn5 transposase. This reaction was incubated for 60 minutes at 23^°^C. Subsequently, the DNA-transposase mixture was dialyzed for 30 minutes to 60 minutes using 0.025 µm dialysis membranes. 0.5 to 1 µL of the dialyzed product was transformed into the desired competent to make the transposon mutagenesis library. The product could also be stored at −20^°^C

### 5. Conjugation

Conjugation experiments were used for constructing the transposon mutagenesis library and the Hfr. Overnight cultures were started for the donor and the recipient strains in 5 mL LB with their respective antibiotics to select for the selection markers and plasmids. In the cases where the recipient was a library, we started 100 mL overnight culture in LB using 500 µL of the library glycerol stock. For the donors in both cases, the media was supplemented with 0.3 mM diaminopimelic acid (DAP) as the strains were auxotroph for DAP. On the subsequent day, fresh 50 mL cultures were launched for the donors and recipients in LB and their respective antibiotics for selection. While the cells grew, LB plates supplemented with DAP were prewarmed at 30^°^C. When the donors and recipients reached an OD of 0.5-0.6, they were centrifuged at 8,600 g for 3 minutes. The cells were washed with 10 mL physiological serum and centrifuged at 8,600 g for 3 minutes at 4^°^C. The washing step was repeated twice. The cells were finally resuspended in 1 mL of physiological serum. Subsequently, the OD was normalized by diluting 10 µL of culture in 990 µL of fresh medium in a spectrophotometer cuvette. The OD600 was verified on the spectrophotometer and the volume of media to add was calculated using the following equation medium to add in µl = (OD0 × V0)/(ODF)– V0, where OD0 is the current OD600 value, V0 is the current volume that the cells are suspended in and ODF is the desired OD600 value. The final OD of cells should be between 15-25. Donors and recipients were combined in a 4:1 ratio by mixing 80 µL + 20 µL of cells. The combined cells were thoroughly mixed and pipetted onto a prewarmed agar plate without antibiotics as two 20-µL spots and si× 10-µL spots as follows. All agar plates were transferred to 37°C for overnight incubation. The cells were removed from each plate by rinsing 750 µl of fresh medium over the plate multiple times. The cells were removed from each plate and the aspirated suspension was placed in a microcentrifuge tube. The cells were centrifuged at 13,500g for 1 min at room temperature to pellet the cells. The cells were serially diluted up to 10^−4^ dilutions and the dilutions were plated on LB agar plates with the required selection antibiotics. The conjugation to construct Hfr using the CRISPR-cas9-recombineering followed the same protocol except that the cells were induced at 42°C after they reached the OD of 0.5-0.6 and placed on ice right after. The conjugation between Hfr donors and recipients followed the same protocol as well, except the cells were plated mixed on LB only as neither the donor nor the recipient had DAP dependence. Additionally, instead of an overnight incubation, we performed the incubation for three hours.

### 6. Calculation of Conjugation Efficiency

The results were presented as colony-forming units per millilitre culture (CFU/mL) relative to the donor. Conjugation efficiencies were calculated by dividing CFUs/mL of the trans-conjugant cells to the CFU/mL of the recipient without selecting the transconjugants. The Hfr frequency was calculated as the number of transconjugants per donor.

### 7. Anti-bacterial activity assay

Our protocol was adapted from Diniz et al. (44) screened strains (HS and 536) are targeting K12::*ZeoR* and are not able to grow on Zeocin. The positive control strain is IAI14 (from O. Clermont’s IAI collection), a toxins-producing strain against K12. The negative control is K12 mixed with K12::*ZeoR.* After OD normalization, the strains are mixed at a 5:1 ratio (potential killer vs K12::*ZeoR*). Plating was done on LBA Zeocin, by serial dilution to quantitatively measure bacterial survival. Biological replicates were performed.

### 8. Statistics and reproducibility

All statistical analyses were performed in R (R Core Team, 2022).

Tests used are indicated in the text or figures’ legend. All data are representative of at least two biological replicates. Where two groups were being compared, a Wilcoxon test was used. p-values below 0.05 were considered significant.

### 9. Genome Sequencing

WGS was performed using Illumina technology. DNA samples were extracted using the Genomic DNA NucleoMag 96 tissue kit from Macherey-Nagel. The whole genome sequencing libraries were prepared and indexed using the Illumina DNA Prep Tagmentation kit and IDT for Illumina DNA/RNA UD Indexes kit A/B/C/D. The paired-end sequencing was performed with Illumina MiniSeq or NextSeq 500/550 to a read length of 2 by 150 bp. The genomes were sequenced at an average depth of 100× for the pooled libraries and 10X for individual clones.

### 10. Bioinformatics

To detect the fraction of the genome that was recombined, reads from the genome of a recombinant clone or a pool of recombinants were mapped simultaneously against the donor and recipient genome with the Burrows-Wheeler mapper – bwa either manually or automatically using a Python script. (45). The recipient genome included the plasmid. Reads were then sorted into three categories using a Perl script or a equivalent python script. The first category was Donor-specific reads which refers to reads matching the Donor genome specifically, that is to say, either no match is found in the Recipient genome or the match on the Recipient genome has a lower score. Similarly, the second category was recipient-specific reads. The third category referred to reads that matched with equal scores to both Donor and Recipient genomes. For the specific reads, when existing, the position of the match in the alternative genome was recorded. Then using an R program, the positions of the reads specific to either the Donor or Recipient genome that had a match in the alternative genome were sorted according to their position in the Recipient genome and associated a Recipient (0) or Donor (1) score. For the genome of individual clones, this table was used to feed a Hidden Markov Model (HMM) with three states: Recipient (R), Donor (D) or a mixture of both (M). The objective of the HMM was to delimit the fraction of the genome that belongs to each of these states. Each state is assigned an emission probability that corresponds to the probability of generating a read with a 1 or a 0 signature along the genome, and a probability of transition between states that was assigned at a very low level: 10^−7^. D state generates 1 with a probability of 0.9, and 0 with 0.1, while R generates 0 with a probability of 0.9 and 1 with 0.1. M generates 0 and 1 with an equal probability of 0.5. It corresponds to a state where the recombination has not been resolved in the colony. Using package HMM (Himmelmann, R package version 2010) and the verbatim algorithm we could recover the recombined fragment. For 536, the analysis was made slightly more complex due to a poor reference genome that had mutations relative to the genome of our strain that could lead to spurious signature of recombination. To avoid these fake recombinations associated with the presence of one or two mutations, recombined fragments with a size lower than 600 bp were discarded as well as recombination detected using the reads produced from our laboratory version of 536 reference genome (46). For the pooled samples, our analysis in R focused on plotting the log ratio of reads Donor, Recipient along the genome.

### 11. Molecular Biology

To look at the depth of genomic analysis, primers were designed upstream and downstream of the detected locus under selection. Primers were designed and then tested for complementarity in Primer-BLAST (Supplementary information). A first PCR reaction was realized to verify the presence or absence of fragments of interest in the amplified region with an expected size of 5 kb. The obtained fragments were sent for Sanger sequencing using another set of primers to amplify < 2 kb regions. From the sequencing results, the sequences were mapped to the reference genomes and the junction region marker/chromosome was identified for each of the samples. The genomic position was compared to the predictive position (maximum of the log ratio donor recipient) to determine the position of the locus.

## Supporting information

**Supplementary Figure 1.**
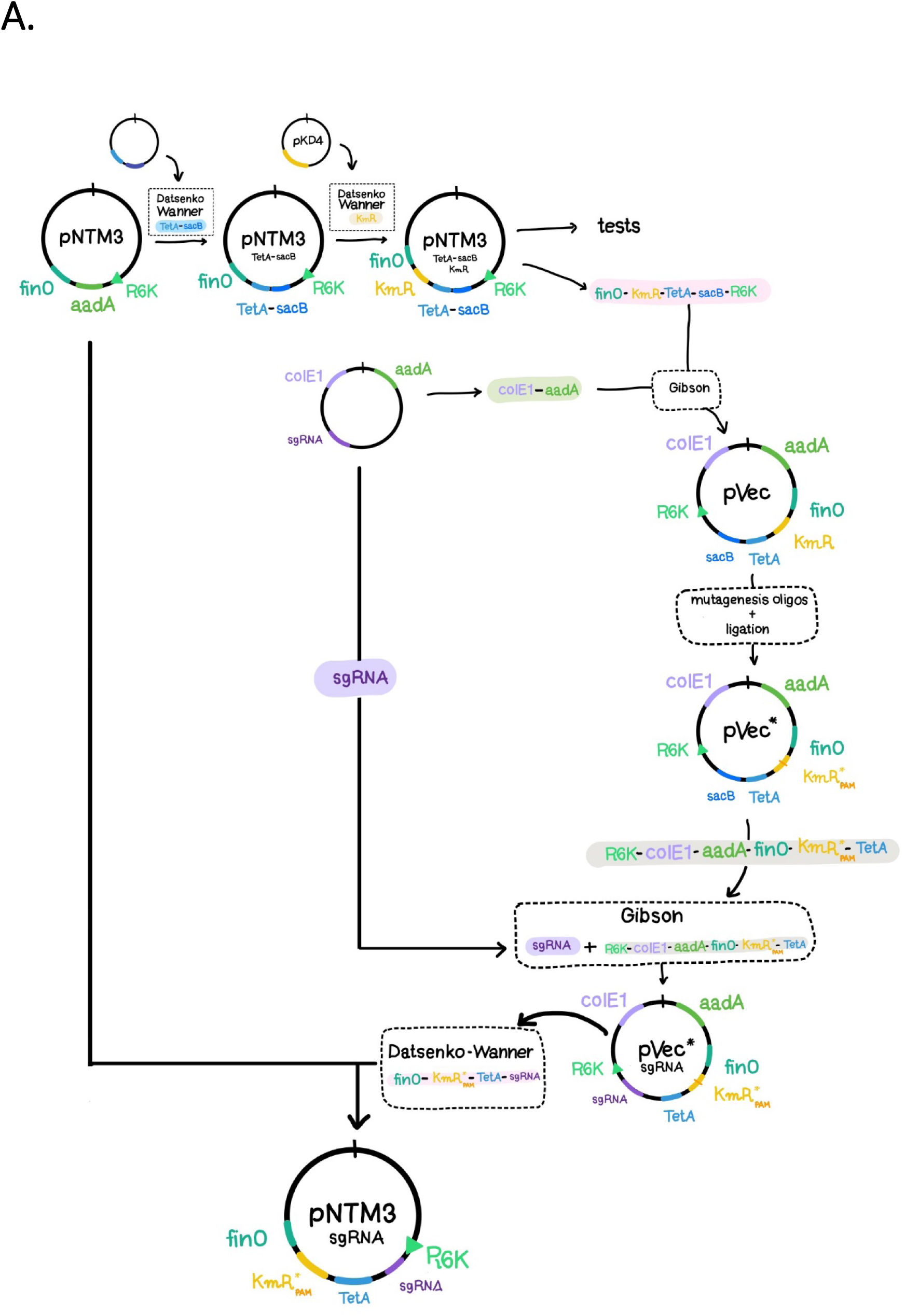
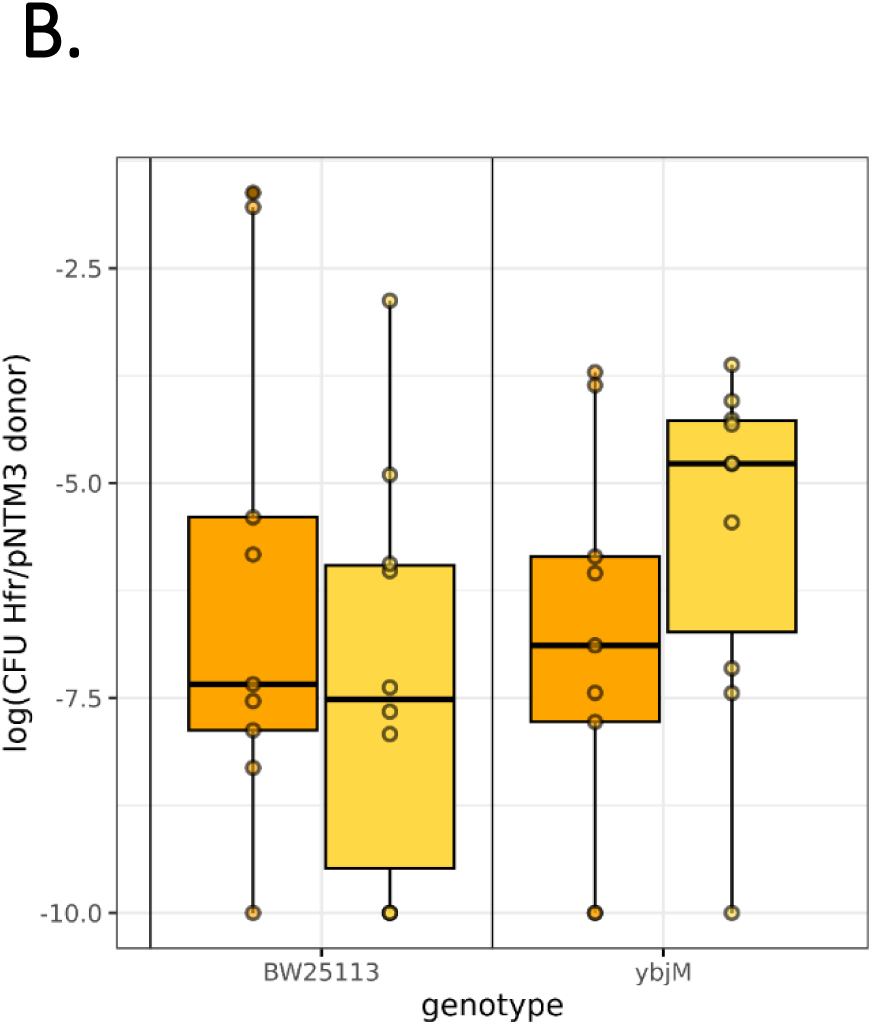
Design of the conjugative plasmid and efficiency of the Cas9-Lambda Red system. A) Construction of the plasmid pNTM3_sgRNA. B) The impact of Cas9-Lambda-red system on plasmid insertion is presented through box plots representing the Hfr frequency measured as ratio of CFU per mL of trans-conjugant recombinants (selected on LB + tetracycline) to CFU per mL of donors (selected on LB + DAP + tetracycline), in cells without (orange) and with the sgRNA (yellow).

**Supplementary Figure 2.**
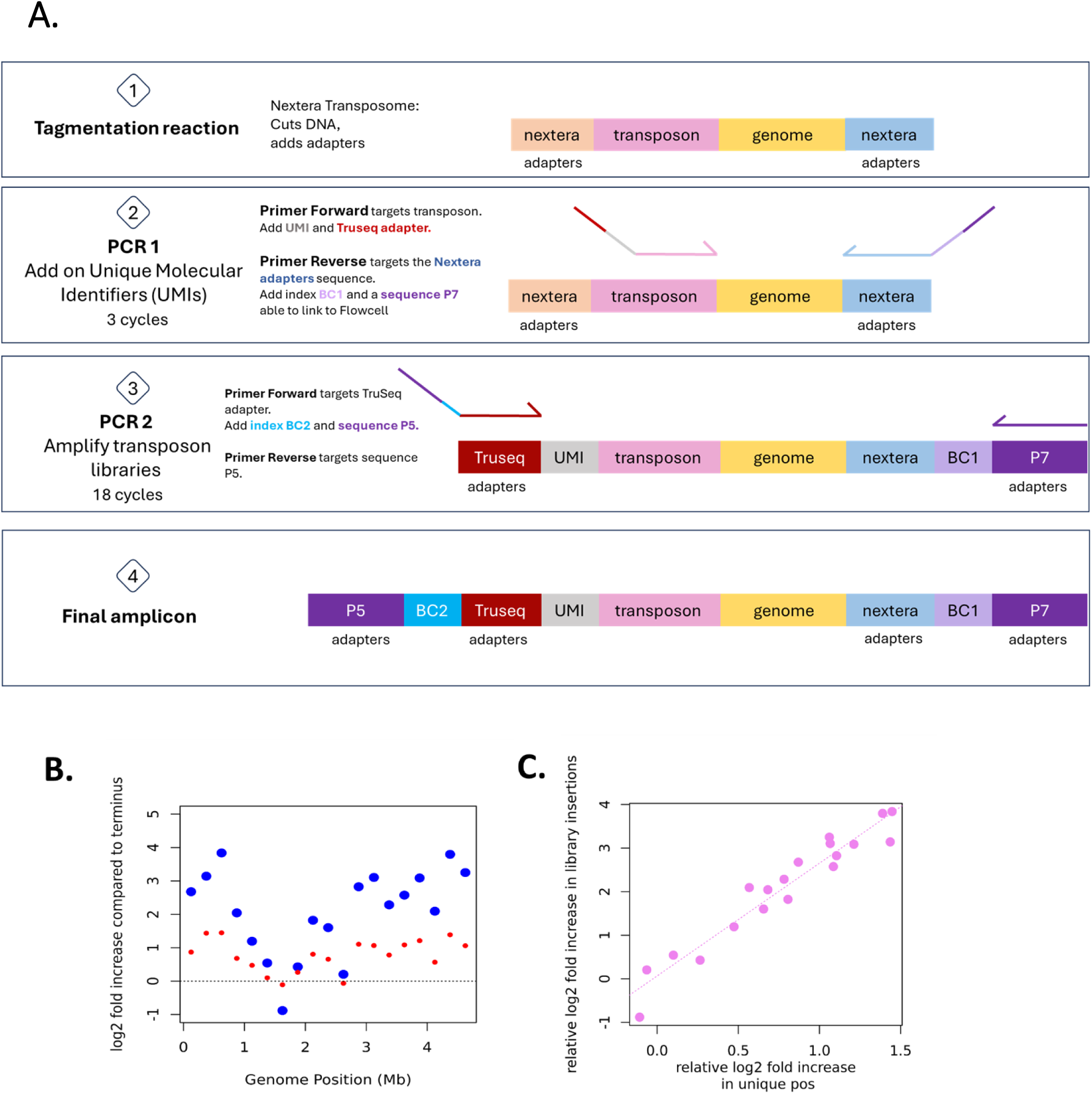
Transposon sequencing of the Hfr donor libraries. A) Methodology for transposon sequencing. B) Transposon Insertion Density and log2 fold increase of the coverage across the chromosome relative to the macrodomain Ter. C) Bias in Coverage of Transposon Insertion Sites across the chromosome.

**Supplementary Figure 3.**
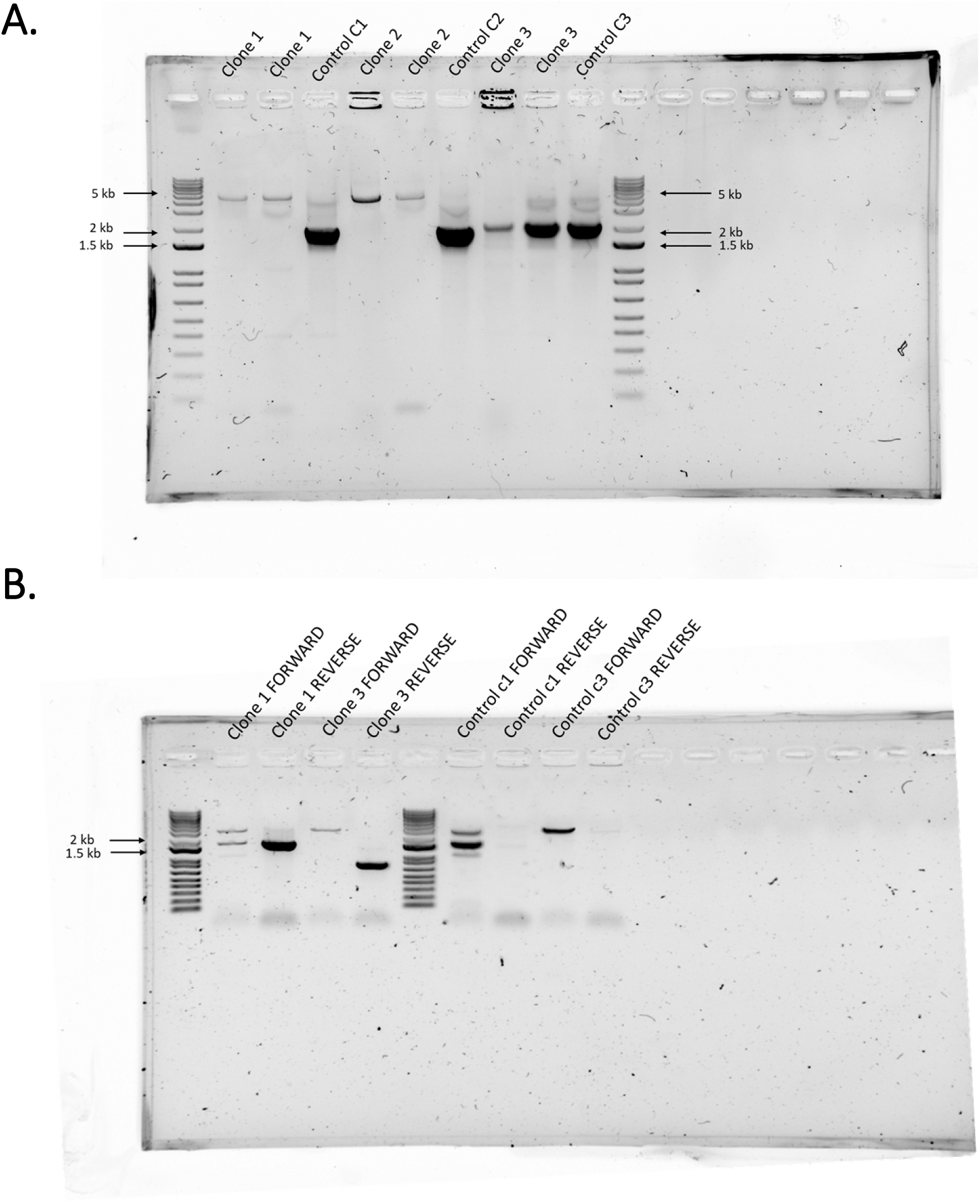
Agarose gel of PCR amplified Tse2 cassette integrated into recombinants. A) Agarose gel of the PCR amplified products to verify the insertion of the 5kb cassette. 3 REL606 clones containing the cassette Tse2 (length ∼ 3kb) were studied, each performed in replicate, using REL606 ancestral strain as a control. The set of primers were indicated on the Supp Material sheet (L_pool_F/ L_pool_R for clone 1 and control c1, M_pool_F/ M_pool_R for clone 2 and control c2, N_pool_F/ N_pool_R for clone 3 and control c3). The presence of a black band at 5 kb indicates the presence of the cassette and at 2 kb that it is absent (targeted region without insertion). For clone 3, no DNA was amplified. New primers were designed to identify a larger region before and after the locus to confirmed the presence of the cassette within a 1.5 kb interval precision. (N_F_before/N_R_before and N_F_after/N_R_after). B) Agarose gel of a niche PCR amplified products to verify the insertion for clone 3 and the sense of insertion of the cassette for clone 1 and 3 and obtained smaller DNA fragments for sequencing. Two clones were studied located on the left and the controls on the right. Two sets of primers were tested: one for each possible sense of insertion. DNA was sent for sequencing for the set of primers used for the Reverse (R), confirming the cassette insertion by the presence of a band for both samples and its absence in the control. For Clone 1 FORWARD/ Control c1 FORWARD: the primers L_pool_F/Tse2_F. For Clone 1 REVERSE/ Control c1 REVERSE: the primers L_pool_R/Tse2_R. For Clone 3 FORWARD/ Control c3 FORWARD: the primers N_F_before/Tse2_F. For Clone 3 REVERSE/ Control c3 REVERSE: the primers N_F_before/Tse2_R.

**Supplementary Figure 4.**
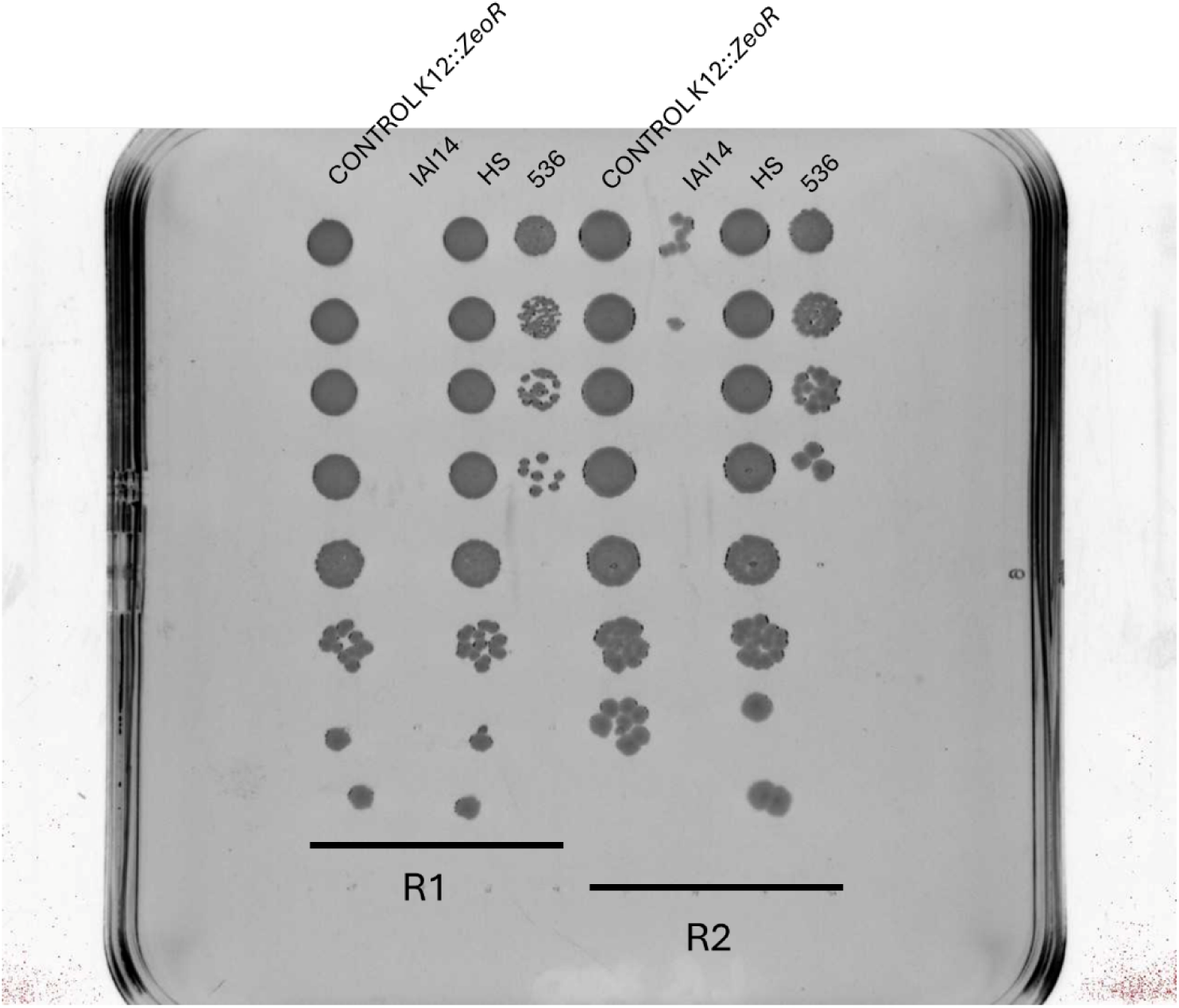
Anti-bacterial activity assay. Serial dilution of anti-bacterial activity assay with K12::*ZeoR* as control and strain of interest. The screened strains are from left to right: control (K12::*ZeoR* versus K12::*ZeoR)*, IAI14, HS, 536, control, IAI14, HS, 536.

**File S1. Supporting Information.** Page 1: strains, plasmids, primers. Page 2 to 9 : Supplementary tables and material (resp. for figures 2B, 3B, 4B, 4C, S1, S2, S3 and S4).

## Acknowledgements

We thank Isabelle Rosinski-Chupin for her critical reading of the manuscript and Celia Rizoug for her contribution to experimental work. We thank Olivier Clermont for his IAI strain collection.

## Funding

This work was supported by ANR GeWiEp (ANR-18-CE35-0005), ANR EcoRecEp (ANR-23-CE35-0006) received by O.T. J.B. received a fellowship from the INCEPTION (ANR-16-CONV-0005) PhD program at Institut Pasteur.

## Notes

### Competing Interest Statement

The authors have declared no competing interest.

